# Predicting cell-cycle expressed genes identifies canonical and non-canonical regulators of time-specific expression in *Saccharomyces cerevisiae*

**DOI:** 10.1101/387050

**Authors:** Nicholas L Panchy, John P. Lloyd, Shin-Han Shiu

## Abstract

The collection all TFs, target genes and their interactions in an organism form a gene regulatory network (GRN), which underly complex patterns of transcription even in unicellular species. However, identifying which interactions regulate expression in a specific temporal context remains a challenging task. With multiple experimental and computational approaches to characterize GRNs, we predicted general and phase-specific cell-cycle expression in *Saccharomyces cerevisiae* using four regulatory data sets: chromatin immunoprecipitation (ChIP), TF deletion data (Deletion), protein binding microarrays (PBMs), and position weight matrices (PWMs). Our results indicate that the source of regulatory interaction information significantly impacts our ability to predict cell-cycle expression where the best model was constructed by combining selected TF features from ChIP and Deletion data as well as TF-TF interaction features in the form of feed-forward loops. The TFs that were the best predictors of cell-cycle expression were enriched for known cell-cycle regulators but also include regulators not implicated in cell-cycle regulation previously. In addition, ChIP and Deletion datasets led to the identification different subsets of TFs important for predicting cell-cycle expression. Finally, analysis of important TF-TF interaction features suggests that the GRN regulating cell cycle expression is highly interconnected and clustered around four groups of genes, two of which represent known cell-cycle regulatory complexes, while the other two contain TFs that are not known cell-cycle regulators (Ste12-Tex1 and Rap1-Hap1-Msn4), but are nonetheless important to regulating the timing of expression. Thus, not only do our models accurately reflect what is known about the regulation of the *S. cerevisiae* cell cycle, they can be used to discover regulatory factors which play a role in controlling expression during the cell cycle as well as other contexts with discrete temporal patterns of expression.

## Introduction

Biological processes, from the replication of single cells (Spellman et al., 1998) to the development of multicellular organisms (Tomancak et al., 2002), are dependent on spatially and temporally specific patterns of gene expression. A gene’s pattern of expression is defined based on the magnitude of expression under a defined set of circumstances, such as a particular environment (Zou et al., 2011; Uygun et al., 2017), anatomical structure (Segal et al., 2008; Chikina et al., 2009), development process (Busser et al. 2012), diurnal cycle (Beer and Tavazoie, 2004; Panchy et al., 2014) or a combination of the above (Uygun et al., 2017). These complex expression patterns are, in a large part, the consequence of regulation during the initiation of transcription. Initiation of transcription primarily depends on transcription factors (TFs) binding to *cis-*regulatory elements (CREs), and other co-regulators to further promote or repress the recruitment of RNA-Polymerase (Juven-Gershon et al., 2008; Lelli et al., 2012; Spitz and Furlong, 2012). This process is influenced by the chromatin state around the promoter and CREs (Miller and Widom, 2003; Benveniste et al., 2014; Li et al., 2015a). In this intricate web of transcriptional initiation regulatory mechanism, TFs play a central role. In addition to CREs and co-regulators, TFs interact with other TFs both cooperatively (Kazemian et al., 2013; Jolma et al., 2015) or competitively (Miller and Widom, 2003). In addition, a TF can regulate the transcription of other TFs. The sum total of TF-target gene and TF-TF interactions regulating transcription in an organism is referred to as a gene regulatory network (GRN) (Macneil and Walhout, 2011).

The connections between components in the GRN are central to the control of gene expression. Thus, knowledge of GRN can be used to model gene expression patterns. Without including regulatory interaction information, CREs have been used to assign genes in *Saccharomyces cerevisiae* into broad co-expression modules (Chikina et al., 2009), to identify enhancer regions involved in myogenesis in *Drosophila* (Busser et al. 2012), classify whether a gene will be stress responsive or not in *Arabidopsis thaliana* (Zou et al., 2011), and predict the timing of diel expression in *Chlamydomonas reinhardtii* (Panchy et al., 2014). Thus far, these studies using CREs to predict expression patterns have had mixed success. Particularly, it remains challenging to predict the timing of expression, such as the diel cycle (Panchy et al., 2014). Beyond using only predicted CREs, inclusion of TF-target interactions has been shown to improve the performance of gene expression prediction in spatially specific stress responsive transcription (Uygun et al. 2017). It is expected that the inclusion of regulatory interaction data will improve the classification of genes expressed in a cyclic, time specific fashion but this expectation remains untested.

The cell cycle of budding yeast, *S. cerevisiae* (reviewed in Bahler 2005) is an ideal system for studying the regulation of cyclic expression patterns. This is because cell cycle gene expression has been extensively characterized in *S. cerevisiae* (Price et al. 1991; Spellman et al. 1998). In addition, transcriptional regulation plays a key role in the cell cycle expression control (Futcher 2002, Breeden 2003). Furthermore, there are multiple data sets defining TF-target interactions in *S. cerevisiae* on a genome-wide scale (Harbison et al. 2004, C. Zhu et al. 2009, Reimand et al. 2010, de Boer and Hughes 2012). These approaches include *in vivo* binding assays, e.g. Chromatin Immuno-Precipitation (ChIP) (Buck and Lieb, 2004; Furey, 2012), *in vitro* binding assays such as protein binding microarrays (PBM) (Bulyk, 2007; Berger and Bulyk, 2009), and comparisons of TF deletion mutants with wildtype controls (Reimand et al., 2010). In this study, we address the central question of how well existing TF-target interaction data can be used to predict gene expression using machine learning algorithms for each cell cycle phase. We also investigated whether performance could be improved by including TF-TF interactions in the form of incoherent feedforward loops as model inputs, applying feature selection algorithms, and by combining interactions from different datasets in a single predictor. Finally, we used the most important TF-TF interactions from our models to construct GRNs that reveal networks of TFs with both known and novel interactions central to controlling the timing of cell cycle expression.

## Results and Discussion

### Comparing TF-target interactions from multiple regulatory data sets

Although there is a single GRN which describes transcriptional regulation in an organism, different approaches to defining regulatory interactions may result in differences in inferred GRNs. Here, TF-target interactions in *S. cerevisiae* were defined based on: (1) ChIP- chip experiments (ChIP), (2) changes in expression in deletion mutants (Deletion), (3) TF PWMs (PWM1), (4) a subset of PWMs curated by experts (PWM2), and (5) PBM experiments (PBM; **Table 1, Methods, Files S1-S5**). The number of TF-target interactions in the *S. cerevisiae* GRNs ranges from 16,602 in the ChIP-Chip data set to 78,095 in the PWM1 data set. This ~5-fold difference in the number of interactions identified is driven by differences in the average number of interactions per TF, which ranges from 105.6 in the ChIP GRN to 558.8 in the PBM GRN (**Table 1**). For this reason, even though most TFs were present in >1 data sets (**Figure 1A**), the numbers of interactions per TF are only weakly correlated between data sets (e.g. between ChIP and Deletion, Pearson’s correlation coefficient (PCC) = 0.09; ChIP and PWM, PCC = 0.11; and Deletion and PWM, PCC=0.046). In fact, for 80.5% for TF, a majority of their TF-target interactions were unique to a single data set (**Figure 1B**), indicating that, in spite of relatively similar coverage TF and their target genes, these data set provide distinct characterizations of the *S. cerevisiae* GRN.

**Table 1.** Size and origin of GRNs defined using each data set.

| Data Set | TF | Target genes | # of interactions | Source |
| --- | --- | --- | --- | --- |
| ChIP | 152 | 4701 | 16,062 | ScerTF |
| Deletion | 151 | 5256 | 26,757 | ScerTF |
| PWM1 | 230 | 6536 | 78,095 | YeTFaSCO |
| PWM2 | 104 | 4740 | 9726 | YeTFaSCO |
| PBM | 81 | 4922 | 45,264 | Zhu et al. (2009) |

**Figure 1.**
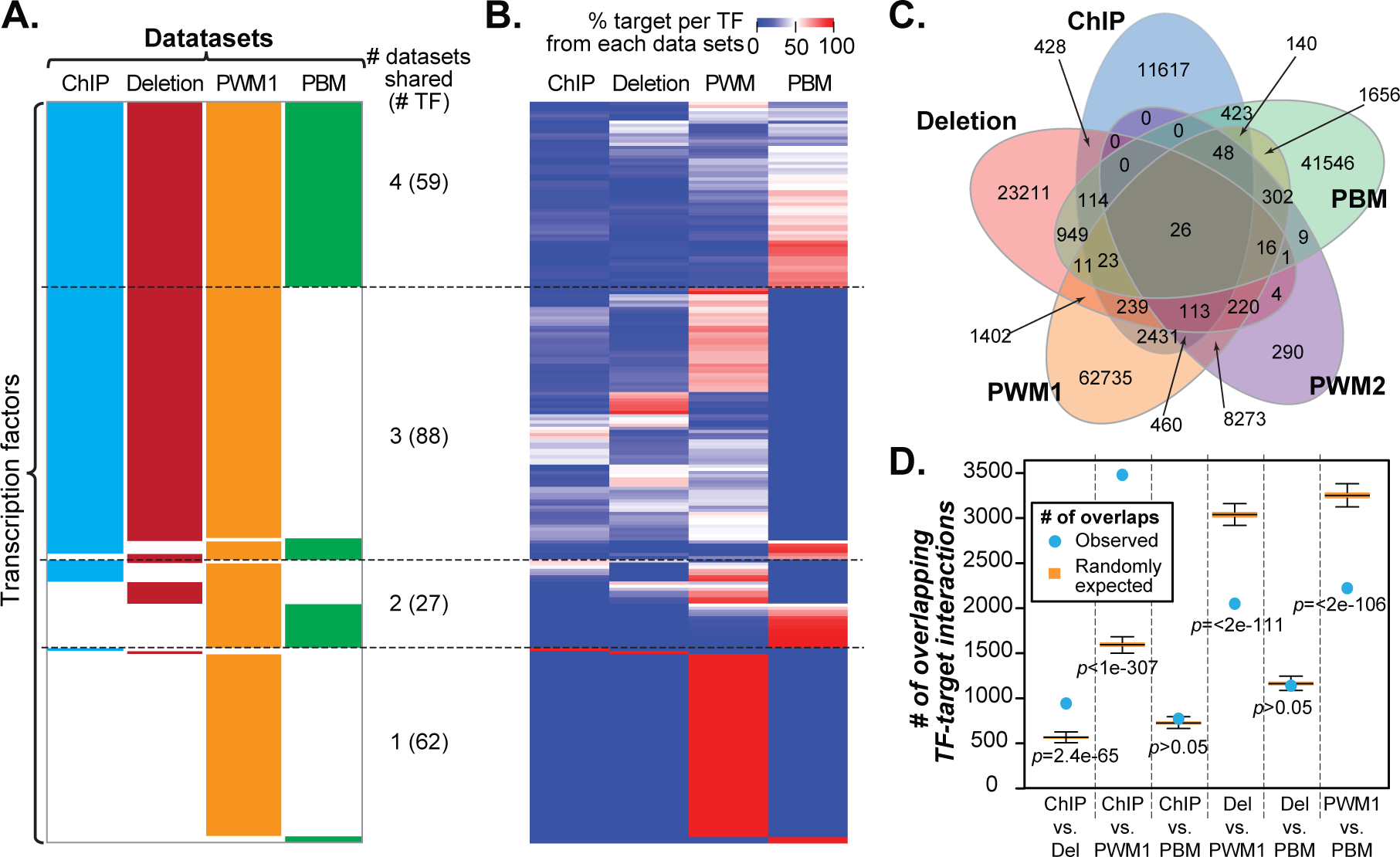
Overlap of TF and interactions between data sets. **(A)** The coverage of *S. cerevisiae* TFs (rows) in GRNs derived from four data sets (columns). ChIP: Chromatin Immuno-Precipitation. Deletion: knockout mutant expression data. PBM: Protein-Binding Microarray. PWM: Position Weight Matrix. The numbers of TFs shared between datasets or dataset-specific are indicated on the right. **(B)** Percentage of target genes of each *S. cerevisiae* TF (row) belonging to each GRN. Darker red indicates a higher percentage of interactions found within a data set, while darker blue indicates a lower percentage of interactions. TFs are ordered as in **(A)**. **(C)** Venn-diagram of the number of overlapping TF- target interactions from different data sets: ChIP (blue), Deletion (red), PWM1 (orange), PWM2 (purple), PBM (green). **(D)** Expected and observed numbers of overlaps between TF-target interaction data sets. Boxplots of the expected number of overlapping TF-target interactions between each pair of GRNs based on randomly drawing TF-target interactions from the total pool of interactions across all data sets (see **Methods**). Blue filled circles indicate the observed number of overlaps between each pair of GRNs.

This lack of correlation is due to a lack of overlap of specific interactions (i.e. the same TF and target gene) between different data sets, (**Figure 1C**). Of the 156,710 TF-target interactions analyzed, 89.0% were unique to a single data set, with 40.0% of unique interactions belonging to the PWM1 data set. Although the overlaps in TF-target interactions between ChIP and Deletion as well as between ChIP and PWM were significantly higher than when TF targets were chosen at random (*p*=2.4e-65 and *p*<1e-307, respectively, see **Methods**), the overlap coefficients (the size of intersection of two set divided by the size of the smaller set) were only 0.06 and 0.22, respectively. In all other cases, the overlaps were either not significant or significantly lower than random expectation (**Figure 1D**). Taken together, the low degree of overlap between GRNs based on different data sets is expected to impact how expression prediction models would perform. Because it remains an open question which dataset would better predict expression, in subsequent sections, we explored using the five datasets individually or jointly to predict cell-cycle phase specific expression in *S. cerevisiae*.

### Predicting phase-specific expression during *S. cerevisiae* cell-cycle using TF-target interaction information

Cell-cycle expressed genes were defined as genes with sinusoidal expression oscillation over the cell cycle with distinct minima and maxima (Spellman et al. 1998). These genes were clustered into five broad categories by Spellman et al. (1998) consisting of 71-300 genes based on the timing of peak expression that corresponds to the G1, S, S/G2, G2/M, and M/G1 phases of the cell cycle (**Figure 2A**). While it is known that each phase represents a functionally distinct period of the cell-cycle, the extent to which regulatory mechanisms are distinct or shared both within cluster and across all phase clusters has not been modeled using GRN information.

**Figure 2.**
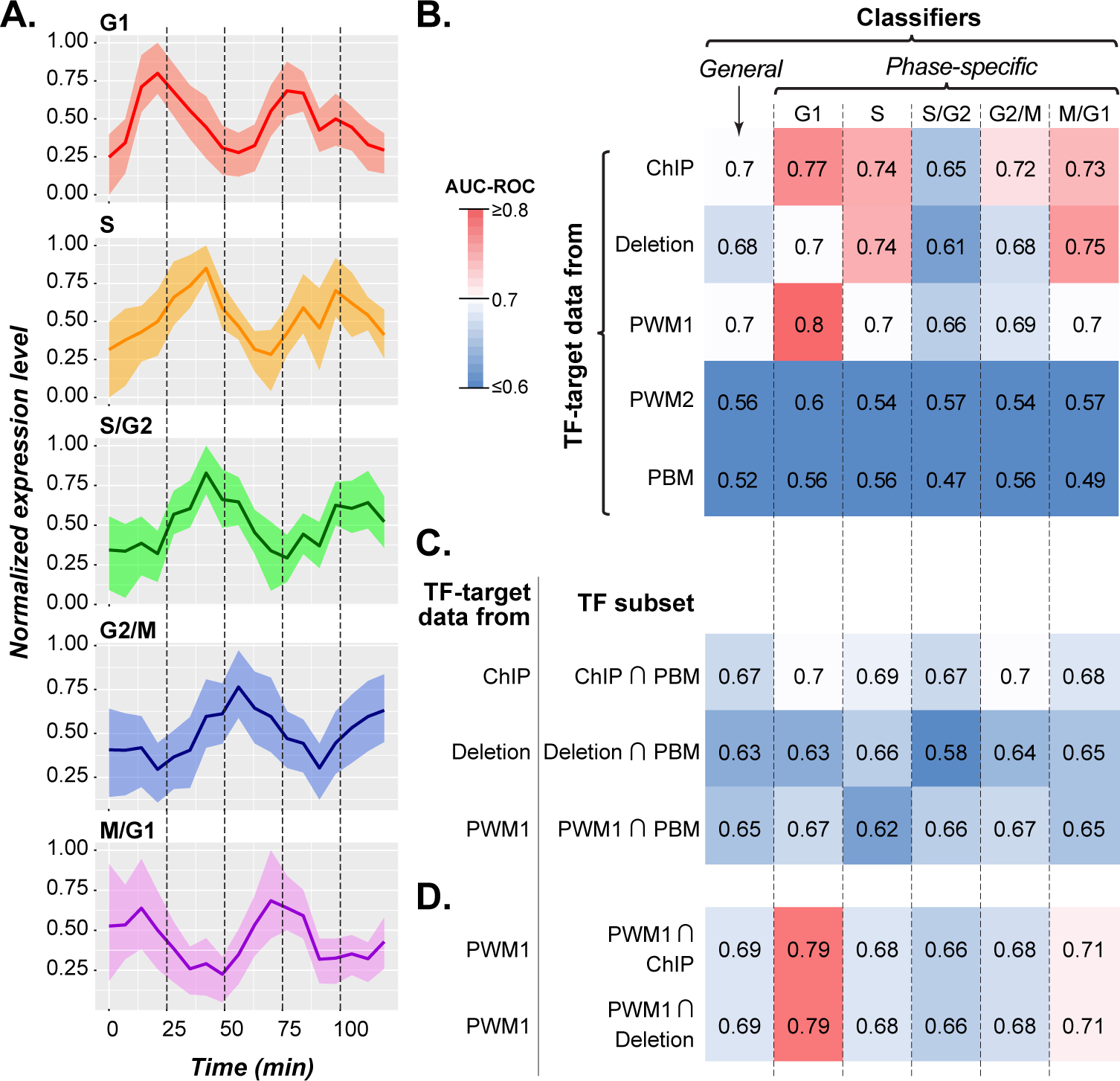
Cell-cycle phase expression and performance of classifiers using TF-interaction data. **(A)** Expression profiles of genes at specific phases of the cell-cycle. The normalized expression levels of genes in each phase of the cell-cycle: G1 (red), S (yellow), S/G2 (green), G2/M (blue), and M/G1 (purple). Time (x-axis) is expressed in minutes and, for the purpose of displaying relative levels of expression over time, the expression (y-axis) of each gene was normalized between 0 and 1. Each figure shows the mean expression of the phase. **(B)** AUC-ROC values of SVM classifiers for predicting whether a gene is cycling in any cell-cycle phases (general) or in a specific phase using TFs and TF-target interactions derived from each data set. The reported AUC-ROC for each classifier is the average AUC-ROC of 100 data subsets l (see **Methods**). Darker red shading indicates an AUC-ROC closer to one while darker blue indicates an AUC-ROC closer to zero. **(C)** Classifiers constructed using the TF-target interactions from the ChIP, Deletion, or PWM1 data, but only for TFs that were also present in PBM data set. **(D)** Classifiers constructed using the TF-target interactions from the PWM1 data, but only for TFs that were also present in ChIP or Deletion data set.

Thus, we used each set of regulatory interactions as features to independently predict whether or not a gene was a cell-cycle gene and, more specifically, if it was expressed during a particular cell-cycle phase using Support Vector Machine (SVM, see **Methods**). Although not all of the regulatory data sets have complete coverage of the *S. cerevisiae* genome, on average the coverage of genes expressed in each phase of cell-cycle was >70% among TF-target datasets (**Table S1**). The performance of the SVM classifier was assessed using the Area Under Curve-Receiver Operating Characteristic (AUC-ROC), which ranges from a value of 0.5 for a random, uninformative classifier to 1.0 for a perfect classifier.

Two types of classifiers were established using TF-target interaction data. The first (general) is for predicting whether a gene is a cell cycle gene in any phase or not. The second type (phase-specific) is for predicting whether a gene belongs to a specific phase cluster. Based on AUC-ROC values, both the data source of TF-target interactions (analysis of variance (AOV), *p*<2e-16) and the phase during the cell cycle (*p<*2e-16) significantly impact prediction performance. Among datasets, the PBM and the expert curated PWM2 dataset have the lowest AUC-ROCs (**Figure 2B**). This poor performance could be because these data sets have the fewest TFs. If we restrict the ChIP, Deletion and full set of PWM (PWM1) data sets to only TF present in the PBM data set, they still perform better than the PBM-based classifier (**Figure 2C**). Hence, the low performance of PBM and the expert PWM must also depend on the interaction inferred for each TF. Conversely, if we take the full set of PWMs (PWM1), which has the most features, and restricts it to only include TFs present in the best performing ChIP or Deletion datasets, performance is unchanged (**Figure 2D**). Therefore, even though a severe reduction of features can impact performance of our classifiers, so long as the most important cell-cycle regulators are covered, performance of the classifier is unaffected.

Overall, our results indicate that both cell-cycle expression in general and timing of cell-cycle expression can be predicted using TF-target interaction data, and ChIP-based interactions alone can be used to predict all phase clusters with an AUC-ROC >0.7, except S/G2 (**Figure 2B**). Nevertheless, there remains room for improvement as our classifiers are far from perfect, particularly for predicting expression in S/G2. One explanation for the difference in performance between phases is that S/G2 bridges the replicative phase (S) and the second growth phase (G2) of the cell-cycle that likely contains a heterogeneous set of genes with diverse functions and regulatory programs. This hypothesis is supported by the fact that S/G2 genes are not significantly over-represented in any Gene Ontology terms (see later sections). Alternatively, it is also possible that TF-target interactions are insufficient to describe the GRN controlling S/G2 expression and higher-order regulatory interactions need to be considered.

### Incorporating feedforward loops for predicting phase-specific expression

Because a gene can be regulated by multiple TFs simultaneously where the regulating TFs may also interact and influence each other’s transcription, our next step was to identify TF-TF-target interactions that may be used to improve phase-specific expression prediction. Here we focused on network motifs, which are patterns of regulatory interactions that are enriched in a biological network and thus theorized to be functionally important (Alon, 2007). In particular, we examined feed-forward loops (FFLs) that consists of a primary TF that regulates a secondary TF and a target gene that is regulated by both the primary and secondary TFs (**Figure 3A**). An FFL is expected to result in peak expression following a delay after the expression of the primary TF is induced (Alon, 2007), and is therefore a potential regulatory mechanism for phase-specific expression in the cell-cycle. We defined FFLs in *S. cerevisiae* using the same regulatory data sets used to identify TF-target interactions and found that significantly more FFLs were present in each of the five GRNs than randomly expected (**Table 2**), indicating FFLs are an overrepresented network motif.

**Table 2.**
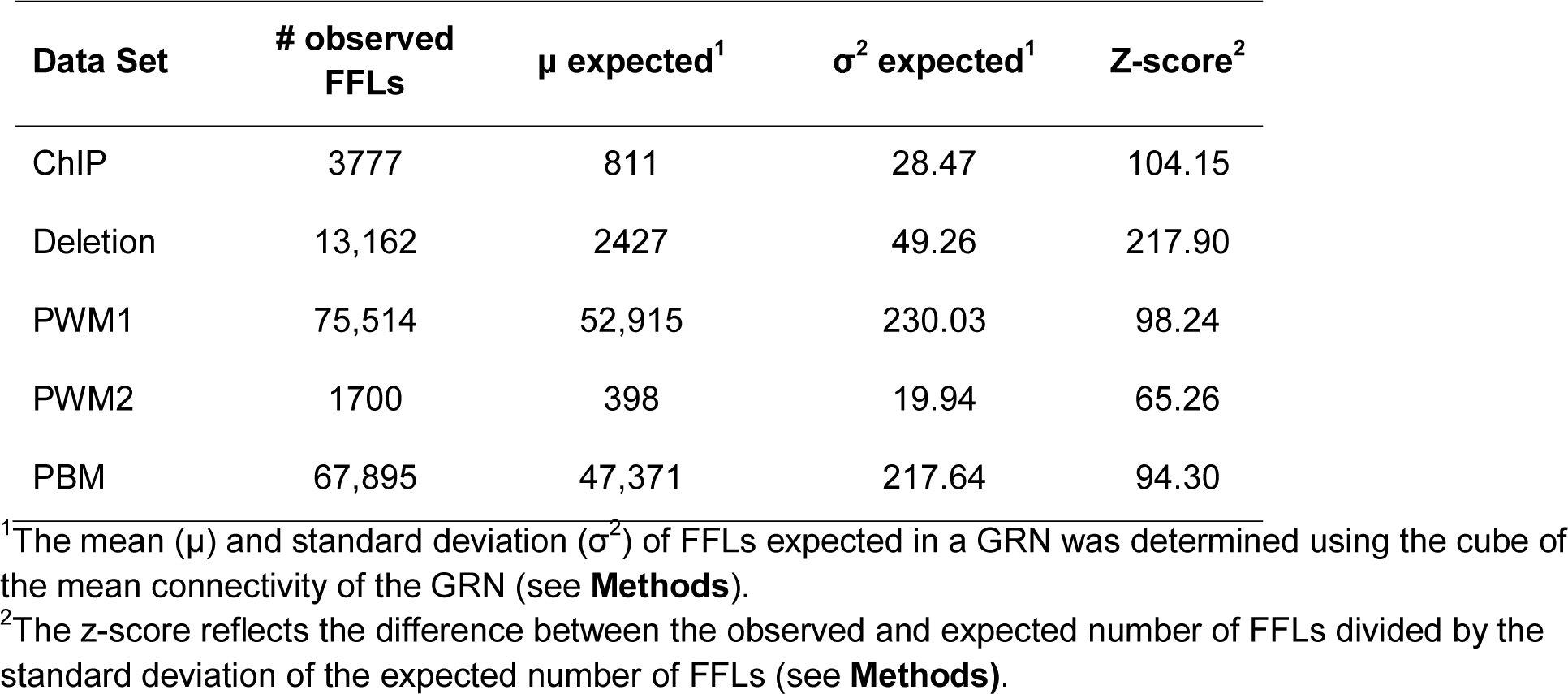
Observed and expected numbers of FFLs in GRNs defined using different data sets.

**Figure 3.**
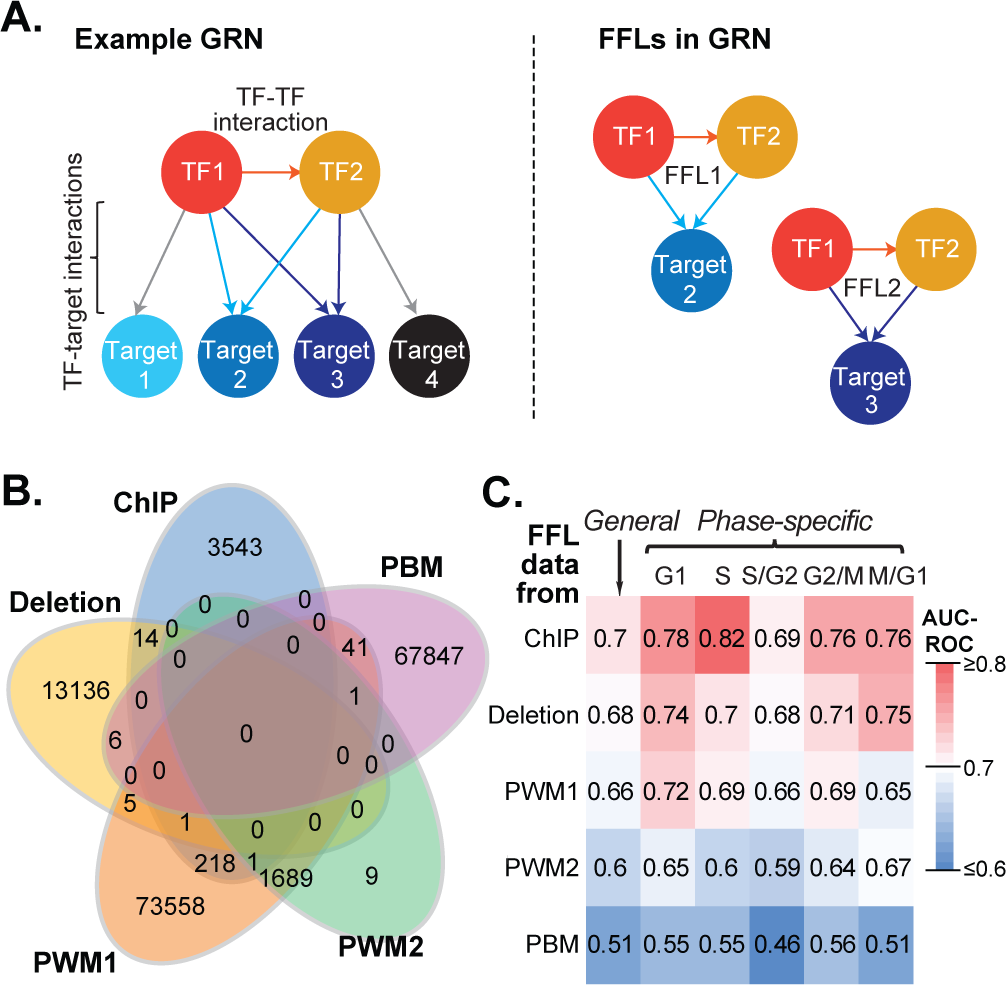
FFL definition and model performance. **(A)** Example Gene Regulatory Network (GRN, left) and feed-forward loops (FFLs, right). The presence of a regulatory interaction between TF1 and TF2 means that any target gene which is co-regulated by both of these TFs is part of a FFL. For example, TF1 and TF2 form a FFL with both Tar2 and Ta3, but not Tar1 or Tar4 because they are not regulated by TF2 and TF1, respectively. **(B)** Correlation plot description. **(C)** Correlation plot description. **(D)** Venn diagram description **(E)** AUC-ROC values for SVM classifiers of each cell-cycle expression gene set (as in Figure 2) using TF-TF interaction information and FFLs derived from each data set. Heatmap coloring scheme is the same as that in **Figure 2**.

We next built models predicting general and phase-specific cell-cycle expression using only regulation by FFLs as features. There was little overlap between data sets ─ 97.6% of FFLs were unique to one data set and no FFL was common to all data sets (**Figure 3B**). Thus, we treated FFLs from each GRN independently in machine learning. Compared to TF-target interactions, fewer cell-cycle genes were part of an FFL, ranging from 19% of all cell-cycle genes in the PWM2 dataset to 90% in PWM1 (**Table S2**). Hence, the models made with FFLs will be relevant to only a subset of cell-cycle expressed genes. Nonetheless, we found the same overall pattern of model performance with FFLs as we did using TF-target data (**Figure 3C**), indicating that FFLs were useful for identifying TF-TF interactions important for cell-cyclic expression regulation. Similar to the TF-target-based models, the best predictions from the FFL- based models were from GRNs derived from ChIP, Deletion, and PWM1. Notably, while the ChIP, Deletion and PWM1 TF-target-based models performed similarly over all phases (**Figure 2B**), predictions using ChIP-based FFLs had the highest AUC-ROC values for all phases of expression (**Figure 3C**).

ChIP FFL models also had higher AUC-ROCs for each phase than those using ChIP-based TF-target interactions. However, if we used ChIP TF-target interactions to predict cell-cycle expression for the same subset of cell cycle genes covered by ChIP FFLs, the performance TF-target based predictions improve better for all phases (**Table S3**). Hence, the improved performance from using FFLs was mainly because the subsets of TFs and cell-cycle gene targets covered by the ChIP FFL provided an effective method for feature selection. Thus, FFLs complement TF-target interactions for the purpose improving model performance. This also suggests that further improvement in cell cycle expression prediction might be achieved by including both TF-target and FFL interactions across data sets.

### Integrating multiple GRNs to improve prediction of cell-cycle expression

To consider both TF-target interactions and FFLs by combining data sets, we focused on interactions identified from the ChIP and Deletion data sets because they contributed to better predictive performance than PBM, PWM1 and PWM2 interactions (**Figure 2B, 3C**. Furthermore, ChIP and Deletion GRNs are expected to be complementary because ChIP identifies direct interactions, while in the Deletion data the interactions can be indirect. To alleviate the concern of overfitting, we first identified which features were important to the performance of the ChIP- based and Deletion-based classifiers. For classifiers based on TF-target interactions, each TF serve as a “feature” for making predictions. For classifiers based on FFLs, a feature is a specific TF-TF combination. The importance of these features was quantified using SVMs weight (see **Methods**) where a positive weight is predictive of cell-cycle expressed genes, while a negatively weighted feature is predictive of non-cell cycle genes. The important features for each phase-specific classifier based on ChIP and Deletion data were defined independently. To determine if model performance can be improved by using only the top features, we defined four feature subsets using two weight thresholds (10th and 25th percentile) with two different signs (positive and negative weights) (see **Methods, Table S4**). This approach allowed us to assess if accurate predictions only require cell-cycle associated (i.e. positive weight) features, or if performance depends on exclusionary (i.e. negative weight) features as well.

First, we assessed the predictive power of each subset of TF-target, FFL, and TF-target/FFL combined features identified using ChIP (**Figure 4A**) or Deletion (**Figure 4B**) data. Generally, the feature subsets consisting of both the top and bottom 25th percentile weights performed best with only one exception, compared to when TF-target and FFL features were considered separately (purple outline, **Figure 4A,B**). When combining TF-target and FFL features, the model performance was not always better compared to using TF-target or FFL data separately. This is to be expected for comparisons to FFL models given that we have previously seen that it is easier to predict the phase of cell-cycle genes regulated by FFLs regardless of what feature set is used (**Table S3**). A notable exception is that models built using the top and bottom 10th percentile of both features was the best predictors of G1 phase (yellow outline, **Figure 4A, B**). In contrast, only M/G1 is predicted better in TF-target models than in models using both TF-target and FFL data, both of which have the same coverage of cell-cycle genes. The results suggest we can achieve equal or improved performance predicting cell-cycle expression via feature selection, so long as both features associated with cell-cycle (positive weight) and non-cell-cycle (negative weight) gene expression are included.

**Figure 4.**
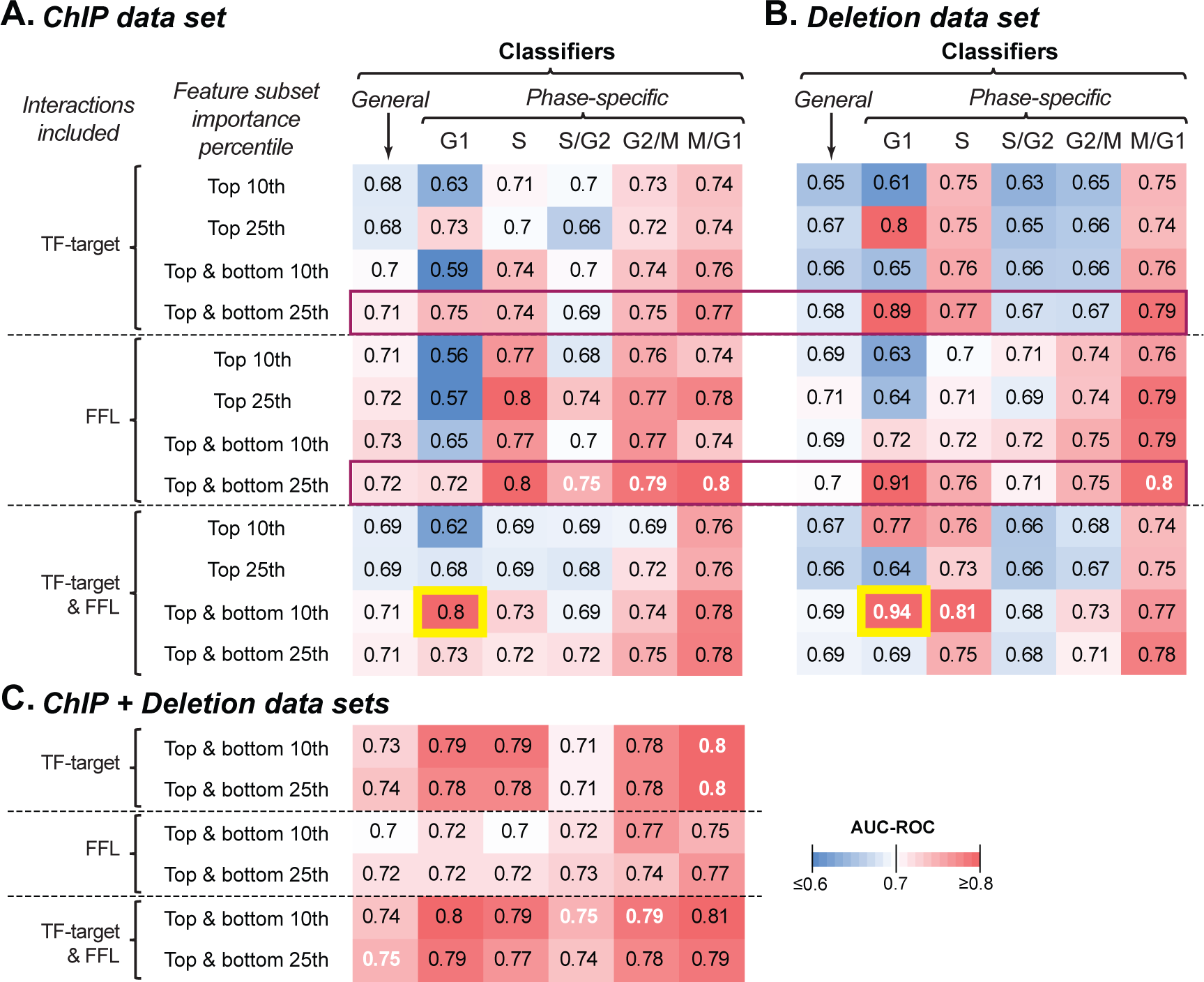
Performance of classifiers using important TF-target and/or FFL features from ChIP, Deletion, and combined data sets. **(A)** AUC-ROC values for models of general cycling or each phase-specific expression set constructed using a subset of ChIP TF-target interactions, FFLs, or both that had the top or bottom 10^th^ and 25^th^ percentile of feature weight (see **Methods**). The reported AUC-ROC for each classifier is the average AUC-ROC of 100 runs (see **Methods**). **(B)** Similar to **(A)** except with Deletion data. **(C)** Similar to **(A)** except with combined ChIP-Chip and Deletion data and only the top and bottom 10^th^ and 25^th^ subsets were used. Purple outline: highlight performance of the top and bottom 25^th^ percentile models. Yellow outline: improved G1-specific expression prediction by combining TF-target and FFL features. White texts: highest AUC-ROC(s) for predicting general cycling genes or genes with peak expression in a specific phase.

Next, we addressed whether combining ChIP and deletion data would lead to models that outperform models using each data type alone. Generally, combining these two datasets (**Figure 4C**) improves or maintains model performance for the general cycling genes and most phase (white texts, **Figure 4**). The ChIP+deletion models were only outperformed by Deletion data set models for G1 and S phase. For general criteria for classifying all phases, the consistency with which classifiers built using both ChIP and Deletion data (**Figure 4C**) outperformed classifiers built with just one data set (**Figure 4A,B**) indicates the power of using complementary experimental data to predict expression. Additionally, these combined models outperform classifiers based on the entirety of any single data set even though they contain fewer total features. While we would expect that this subset of important TFs will consist of known cell-cycle regulators, we also sought to use important TF-target and TF-TF interactions to discover novel TF functions that are associated with cell-cycle regulation.

### Functions of TFs important for predicting cell-cycle expression

In our analysis of the ChIP and Deletion data sets, we found that performance of classifiers using only the most important features is similar to those using the full feature set. Of the 25 TFs that have been annotated as cell-cycle regulators in *S. cerevisiae* (GO:0051726), 20 were covered by the ChIP and Deletion data set (**Table S5**). For ChIP-based classifier, the 10th percentile of the most important TFs from all phases, except M/G1, are enriched for known cell-cycle regulators (**Table 3**). While this pattern of enrichment was not found in Deletion features nor in 25th percentile of features for either data set, 17 of the 25 annotated, cell-cycle regulators are important features for predicting ≥ 1 phase of cell cycle in one or both data sets (**Table S5**). These seemingly contradictory results between enrichment and coverage occur because the set of annotated cell-cycle regulators contains both general and highly phase-specific regulators (**Table S5**).

**Table 3.**
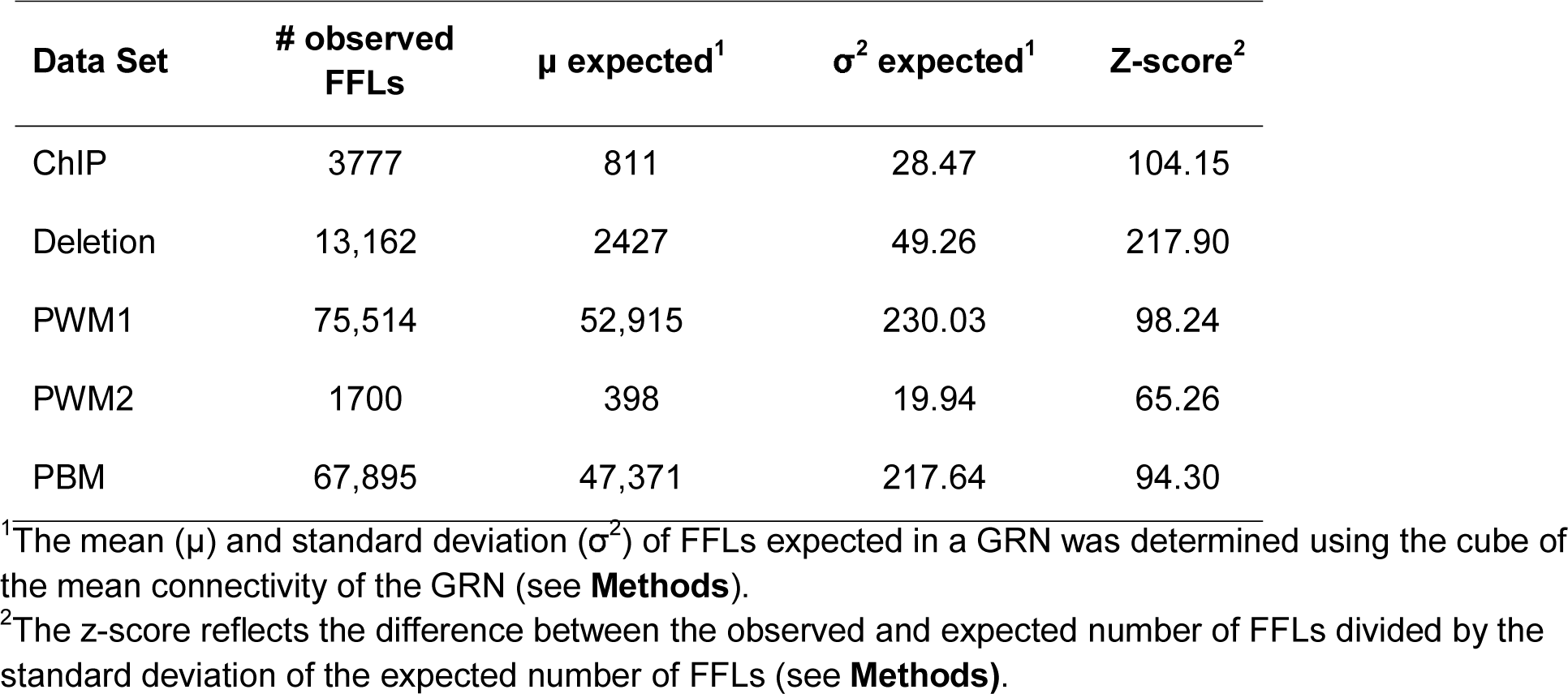
Enrichment *p-*values of known cell-cycle regulators among TF features important to predicting general cell-cycle or phase-specific expression.

While important TFs identified by our classifiers are enriched for annotated cell-cycle regulators, these known regulators still represent the minority of TFs predicted as important. For example, there are 84 TFs with the top 10^th^ percentile weights in the combined model for G1 prediction (**Figure 4**). To better understand the functions of these other important (with high weight) TFs, we looked for enriched GO Terms other than cell-cycle regulation among TFs with the top 10th and 25th percentile weights in classifier predicting general cyclic expression using either the ChIP or the Deletion TF-target data **(Table S6**). We identified 126 over-represented GO terms in total, 94 of which were unique to ChIP-based or Deletion-based classifiers. TFs important in ChIP-based classifiers tend to be enriched in genes involved in the positive regulation of transcription in response to variety of stress conditions (e.g. freezing, genotoxicity, heat, high salinity, reactive oxygen species, and amino acid starvation; **Table S6**). This is consistent with the finding that cell-cycle genes, particularly those involved in the G1-S phase transition, are needed for heat-shock response (Jarolim et al., 2013). In contrast, TFs important to Deletion-based classifiers are enriched in categories relevant to cellular metabolism (e.g. amino acid metabolism, glycolysis, and respiration; **Table S6**), consistent with the view that the metabolic status of the cell determines cell cycle progression (Cai and Tu, 2012).

The distinct functions enriched in TFs important in ChIP and Deletion data supports the hypothesis that the improvement in predictive power from combining feature sets between ChIP and Deletion data was due to the distinct, but complementary characterization of gene regulation in *S. cerevisiae*. These findings also support the notion that a significant number of TFs important for cell-cycle expression predictions are novel cell-cycle regulations.

### Interaction between TFs important for predicting cell-cycle expression

To explore the potential regulatory differences between the ChIP and Deletion datasets, we constructed ChIP and Deletion GRNs by selecting the top 10th percentile of TF features for predicting general cell-cycle expression. The resulting network shows differences in connectivity of GRNs, with only 2 of 15 features in the ChIP being singletons (**Figure 5A**), while 10 of 15 TF are not connected to any other TF in the Deletion network (**Figure 5B**). In addition, only two nodes (MBP1 and SWI4) are shared between these two GRNs (orange outline, **Figure 5A, B**). This connectivity differences likely reflect the nature of the methods in assessing interactions, one direct and the other indirect. The Swi6-Swi4-Mbp1 module, which regulates G1/S phase transition (Iyer et al., 2001; Bean et al., 2005; Wittenberg and Reed, 2005) and part of the Fkh1- Fkh2-Ndd1 module, which regulates S/G2 (Zhu et al., 2000) and G2/M (Koranda et al., 2000) expression, are present in the ChIP but not the Deletion data-based network. We would expect this outcome for the Deletion GRN, as the 10th percentile of important TFs was not enriched for known cell cycle regulations (**Table 3**).

**Figure 5.**
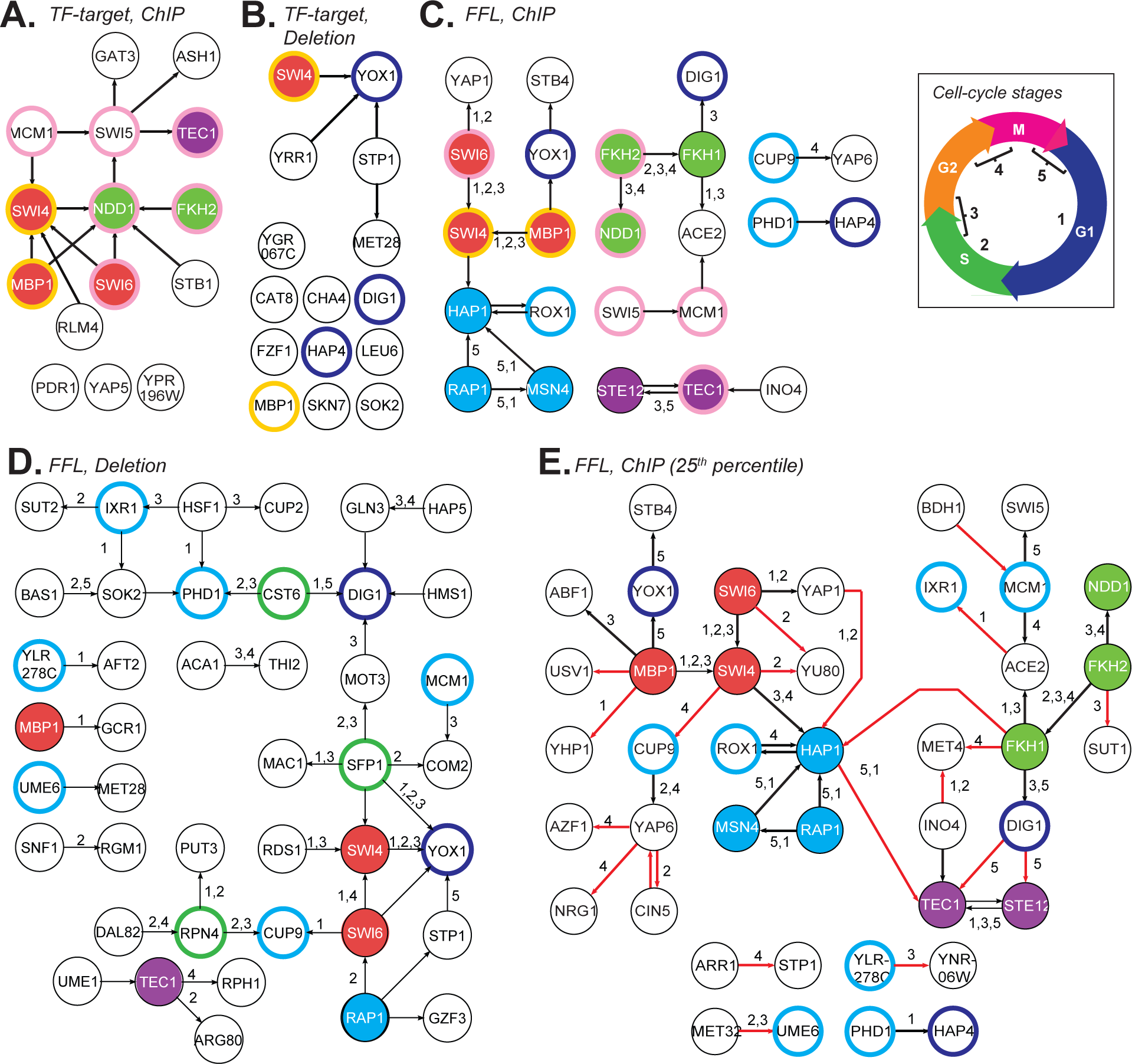
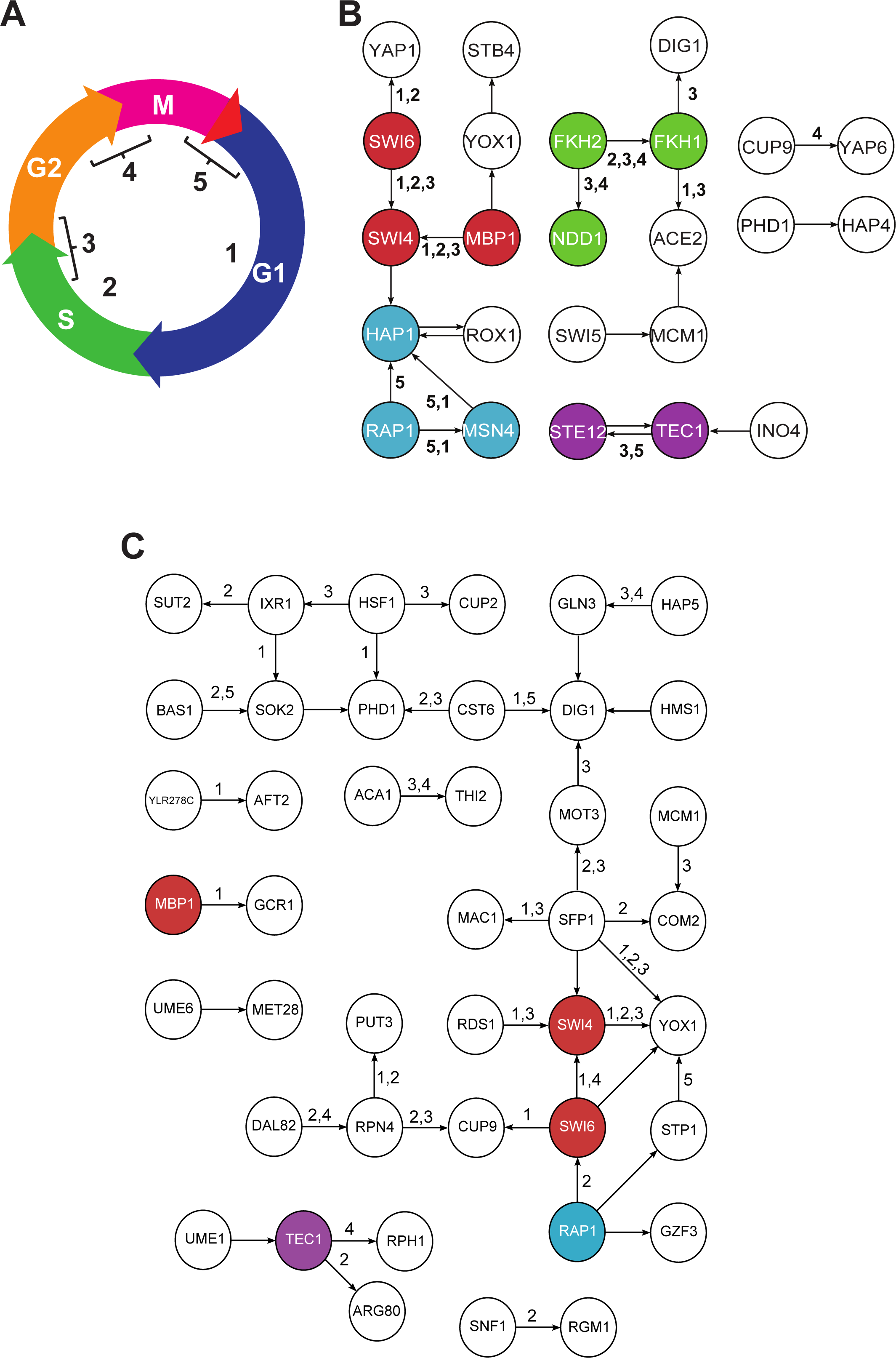
Cell-cycle GRNs based on important TF Features. **(A, B)** The GRNs consisted of TFs with the top 10th percentile weights for predicting all cell- cycle expressed genes using TF-target interactions from ChIP **(A)**, or Deletion **(B)** data. **(C, D)** The GRNs consisted of TFs in FFLs with the top 10^th^ percentile weights for predicting all cell-cycle expressed genes using ChIP (**C)** or deletion **(D)** data. Interactions are further annotated with the phase of cell-cycle expression they are important for predicting (10^th^ percentile of SVM weight in ChIP-Chip models). Insert: Cell-cycle phase 1 = G1, 2 = S, 3 = S/G2, 4 = G2/M, 5 = M/G1. **(E)** The ChIP GRN consisted of TFs in FFLs with the top 25th percentile weights for predicting cell-cycle expressed genes. Red edges: new interactions identified compared to **(C)**. In **(A-E**), node outline colors indicate TFs shared between GRNs in: orange - **(A)** and **(B)**; pink - (A) and **(C)**; blue – **(B)**, **(C)**, **(D)**, and **(E)** (except Hap4); cyan: **(C)**, **(D)**, and **(E)** Filled colors: four modules with TF-TF interactions important for predicting expression in ≥2 phases.

We should also point that while Swi6-Swi4-Mbp1 is present in the ChIP GRN, Fkh1 is missing (**Figure 5A**), suggesting that we may be missing important interactions if we only consider TFs that are individually important. To address this issue, we also built GRNs with the TF-TF interactions in the top 10th percentile of importance ranks from the FFL-based models predicting general cell cycle genes based on ChIP (**Figure 5C**) and Deletion (**Figure 5D**) data. Since these TF-TFs interactions were also used as features in phase-specific predictions, we labeled interactions that were above the 10th percentile of importance for individual phases (edge labels, **Figure 5C, D**). In the GRN based on the ChIP FFL data (**Figure 5C**), 61% interactions were important for predicting ≥1 phases of cell-cycle expression. Furthermore, both Swi6-Swi4-Mbp1 (red) and Fkh1-Fkh2-Ndd1 (green) modules are fully represented in this network and are important for predicting multiple phases of cell cycle expression (**Figure 5C**).

In contrast, using the 10th percentile of TF-TF features based the Deletion data to construct a GRN dataset revealed none of the modules uncovered using the ChIP data (except Swi4 and Swi6, **Figure 5D**). This is most likely because the inferences based on the Deletion dataset were indirect. Nonetheless, the Deletion data allows for the identification of known cell cycle regulators not found in the ChIP network, particular Sfp1 (Xu and Norris, 1998) that also plays roles in regulation of ribosomes in response to stress (Jorgensen et al., 2002; Marion et al., 2004) (green outline, **Figure 5D**). These findings highlight the importance of incorporating TF-TF interaction data, as well as the need to incorporating both ChIP and Deletion datasets.

### Novel regulators of cell-cycle expression

In the GRN based on the ChIP FFL data, two modules that are not annotated as cell-cycle regulators are also identified. The first is the feedback loop between Ste12 and Tec1 important during S/G2 and M/G1 transitions (purple, **Figure 5C**). Ste12 and Tec1 are known to form a complex that shares co-regulators with Swi4 and Mbp1 to promote filamentous growth (van der Felden et al., 2014). The second is the Rap1-Hap1-Msn4 module important for predicting the M/G1 and G1 phases (blue, **Figure 5C**). Rap1 is involved in telomere organization (Guidi et al. 2015; Laporte et al. 2016). Hap1 is an oxygen response regulator (Keng 1992; Ter Linde and Steensma 2002). Msn4 is a general stress response regulator (Martinez-Pastor et al. 1996; Schmitt and McEntee 1996). We should emphasize that none of the TFs in these two modules is annotated as cell-cycle regulators despite their importance in predicting cell-cycle expression. In the Deletion FFL GRN (**Figure 5D**), TFs that are potentially novel cell-cycle regulators can also be identified. Two examples include Rpn4 and Cst6 are responsible for regulating proteolytic stress response (Mannhaupt et al., 1999; Ng et al., 2000; Xie and Varshavsky, 2001) and carbon utilization (Garcia-Gimeno and Struhl, 2000), respectively (green outlines, **Figure 5D**). While these non-canonical regulators do not appear to regulate cell-cycles transitions directly, the types of genes they regulate (metabolism, adhesion/filamentous growth, and stress response) are cell-cycle associated.

Note that, in addition to the identified modules, five TFs (CUP9, DIG1, PHD1, ROX1, and YOX1) are common between the ChIP-based and the Deletion-based TF-TF networks (**Figure 5C, D**). If we are more inclusive and consider the top 25 percentile TF-TF interaction features based on ChIP, four additional TFs are now common (IXR1, MCM1, UME6, YLR-278C; **Figure 5E**). Three of these overlapping TFs (MCM1, UME6, and YOX1) are GO annotated cell-cycle regulators. Of the remaining six, four genes do not directly regulate transition between cell-cycle phases but influence the progression of growth phases, and thus affect the cell-cycle indirectly. DIG1 and ROX1 knockouts delay G1 progression (White et al., 2009), PHD1 interacts with TUP1 (Hanlon et al., 2011), which also results in G1 delay when knocked out (White et al., 2009), and CUP9 regulates PTR2 (Hauser et al., 2001), which causes abnormal G2 progression when over-expressed (Spoko et al., 2006). For the remaining two genes, IXR1 is not known to directly or indirectly affect growth phase progression, but is responsible for aerobic repression the cytochrome c gene COX5B alongside ROX1 (Lambert et al., 1994) and affects the expression of genes involved in DNA replication during hypoxia (Vizoso-Vasquez et al., 2012), but otherwise has not implicated in cell-cycle regulation. YLR-278C lacks any evidence of association with the cell-cycle and its only GO biological progress annotation is “biological progress unknown”. The repeated appearance of YLR-278C in our networks warrants further investigation of the TF as a regulator of the cell-cycle or of cell-cycle related processes.

Overall, these findings demonstrate the utility of the FFL-based classifiers and the need to consider the importance ranks of TF-TF interaction features when predicting gene expression. The GRN constructed from carefully selected TF-TF interactions allow the recovery of regulatory modules which cannot be identified based on TF-target interaction data. Furthermore, GRNs built from the ChIP and Deletion TF-TF interactions both identified interactions important to >1 phases of cell-cycle expression, but the characteristics of these interactions differ. ChIP-based interactions contain modules with known shared functions, while Deletion-based interactions involve central metabolism regulators like Sfp1 and consist of both direct and indirect relationships.

## Conclusions

Predicting the expression of genes from their regulators and regulatory interactions remains a challenging exercise, but one that can be useful for studying how organisms respond to various stimuli and how that response is regulated at the molecular level. Here, we have shown that the problem of predicting complex expression patterns, such as the timing of expression across the cell-cycle, is tractable using a variety of experimental and computational methods of defining TF-target interactions. In spite of painting distinctly different pictures of the *S. cerevisiae* GRN, interactions inferred from ChIP-Chip, Deletion and PWM data sets were useful for predicting genes expressed during the cell cycle and for distinguishing between genes expressed at different phases. In fact, the differences between the ChIP and Deletion sets meant that integrating them into a single model improved the overall accuracy of machine learning predictions. Furthermore, we found that models were improved with the addition of TF-TF interactions in the form of FFLs. Particularly, a subset of the most important TF-TF interactions, combined with a subset of the most important TF-target interactions, led to models that performed better than either the full set of TF-target interactions or FFLs.

By studying the TFs involved in the most important TF-target interactions and FFLs we were able to infer that these interactions play a biologically significant role in regulating the cell-cycle. We found that the 10th percentile of important TFs from every phase except M/G1 were enriched for TFs with cell-cycle annotations. For the M/G1 phase we identified important TF-TF interactions that involve non-canonical cell-cycle regulators, such as the regulatory modules Ste12-Tec1 and Rap1-Msn4-Hap1. The Rap1-Msn4-Hap1 module stands out in that, while these regulators are individually poor predictors of cell-cycle expression, interactions between these TFs are among the best predictors of both cell-cycle expression in general and of the M/G1 and G1 phases in particular. Our GO analysis also indicated that TFs important for predicting cell-cycle expression were enriched for genes associated with metabolism (Cst6), invasive growth (Ste12-Tec1), and stress responses (Rpn4, Rap1-Msn4-Hap1), which was reflected in the network analysis as we found that interactions important for >1 phases of cell-cycle expression were clustered around TFs involved in those processes.

Although our best performing model was based on data with nearly complete coverage of the *S. cerevisiae* TF-DNA interactions, our models do not provide a complete picture of the regulation of cell-cycle expression for at least the following three reasons. First, cell-cycle control involves additional levels of regulation beyond transcription. In particular, kinases and the interaction between kinases and TFs are known to play a key role in regulating the timing of the cell cycle, and FFLs are frequently observed in this TF-kinase network (Csikász-Nagy et al., 2009). Second, better characterization of TF binding sites will also help provide more accurate representation of the GRN regulating expression timing, such as novel methods of characterizing binding sites that incorporate information about both position and DNA modification (Csikász-Nagy et al., 2009; O’Malley et al., 2016). Third, our approach to understanding interactions between TFs involve FFLs, a relatively simple type of network motifs. More complicated interactions involving >2 TFs could further improve the prediction model.

Nevertheless, the fact we were able to predict certain aspects of cell-cycle expression using only FFLs justifies their use in an expression modeling context. Furthermore, FFLs can be used to compose more complex interactions. For example, negative-feedback loops, which have previously been identified as being involved in the regulation of biological oscillations (Bertoli, Skotheim, and de Bruin 2013; Pett et al. 2016), are composed of two FFL where the primary or secondary TFs are reversed. Our identification of the interactions Ste12 and Tec1 as important to cell-cycle expression is an example of how more complicated regulatory pathways can be captured by using their constituent FFLs.

This work shows that predictive models can provide a framework for identifying both regulators and regulatory interactions controlling temporal gene expression. Understanding the molecular basis of the timing of expression is of interest not only for the cell-cycle, but other important biological processes, such as response to acute stresses like predation and infection, and to cyclical changes in the environment including light and heat, and other cues. Although there remains room for improvement, the approach described here is not limited to the study of expression timing, but can also be applied to any expression pattern with discrete phases.

## Methods

### TF-target interaction data and regulatory cite mapping

Data used to infer TF-target interactions in *S. cerevisiae* were obtained from the following sources: ChIP-Chip (Harbison et al., 2004) and Deletion (Reimand et al., 2010) data were downloaded from ScerTF (http://stormo.wustl.edu/ScerTF/), PWMs (de Boer and Hughes, 2012) and the expert curated subset of these PWMs were downloaded from YetFaSCO (http://yetfasco.ccbr.utoronto.ca/), and PBM binding scores were taken from Zhu et al. (see Supplemental Table 5, (Zhu et al., 2009). For ChIP-Chip and Deletion data, the interaction between TF and their target genes were directly annotated, however, for PWMs and PBMs data we mapped inferred binding sites to the promoters of genes in *S. cerevisiae* downloaded from Yeastract (http://www.yeastract.com/). All position weight matrices were mapped for the PWM data set, however for PBM data we only used the oligonucleotides in the top 10th percentile of scores for every TF. This threshold was determined using a pilot study which found that using the 10th percentile as a cutoff maximized performance of prediction using PBM data. Mapping was done according to the pipeline previously described in Zou et al. (2011) using a threshold mapping p-value of 1e-5 to infer a TF-target interaction.

### Overlap between TF-target interaction data

To evaluate the significance of the overlap in TF-target interactions between different GRNs, we compared the observed number of overlaps to what we expected were the genes regulated by each transcription factor randomized. In detail, for each set of TF-target interactions we replaced the target gene of each interaction with one that was randomly drawn from the total set of target genes across all data sets, such that the number of interactions for each TF were preserved. For each randomization of target gene, the number of overlapping features between each pair of data set was calculated. This process was repeated 1000 time to determine the mean and standard deviation of overlap between each data set expected under this randomization regimen. To determine to degree to which our observation differed from the expectation under this random model, we applied the two-tailed z-test to the differences between the observed number of overlaps and the distribution of overlaps from the randomized trials.

### Expected feed-forward loops in *S. cerevisiae* regulatory networks

FFLs were defined in each set of TF-target interactions as any pair of TFs with a common target genes where a TF-target interaction also existed between one TF (the primary TF) and the other (the secondary TF) which, for clarity, we refer to as a TF-TF interaction. The expected number of FFLs in each data set was determined according to the method described by Uri Alon in “An Introduction to Systems Biology” (Chapter 4, 2007b). Briefly, the expected number of FFLs (N_FFL_) in a randomly arranged GRN is approximated by the cube of the mean connectivity (λ) of the network with a standard deviation equal to the square-root of the mean. Therefore, for each data set we compared the observed number of FFLs to the expected number of FFLs from a network with the same number of connections, but with those connections randomly arranged by defining λ as the number of TF-target interactions divided by the total number of nodes (TFs+target genes) and calculating mean the standard deviation as above.

### Validating FFLs in cell-cycle expression

FFLs were validated in the context of cell-cycle expression by modeling the regulation and expression of genes involved in the FFL using a system of ordinary differential equations:

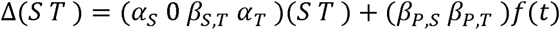

Where S and T are the expression of the secondary TF and target gene respectively, ∝_S_ and ∝_T_ are the decay rates of the secondary TF and target gene respectively, and β_S, T_ indicates the production rate of the target gene dependent on the secondary TF. In the nonhomogeneous term portion of the equation, β_P,S_ and β_P,T_ are the production rate of the secondary TF and target gene, respectively, which depend on the primary TF, while f(t) is the expression of the primary TF over time which is independent of both the secondary TF and the target gene. This system was solved in Maxima (http://maxima.sourceforge.net/index.html). For each FFL, maximum likelihood estimation, implemented using the bbmle package in R (https://cran.r-project.org/web/packages/bbmle/index.html), was used to fit the model parameters to the observed expression of genes during the cell-cycle as defined by Spellman et al. (1998). Each run was initialized using the same set of initial conditions and only FFLs for which a reasonable (∝ < 0, βs > 0), non-initial parameters could be fit were kept. Between 80 and 90% of FFLs in each data set passed this threshold, while only 21% of FFLs built from random TF-TF-target triplets were fit.

### Classifying cell-cycle genes using machine learning

Predicting cell-cycle expression and phase of cell-cycle expression was done using the Support Vector Machine (SVM) algorithm implemented in Weka (Hall et al. 2009). Each expressed gene was treated as separate instance. The features were the presence or absence of TF-target and/or TF-TF interactions in FFLs defined with each of five regulatory datasets (ChIP-Chip, Deletion, PWM, Expert-PWM, and PBM). For the general model, two classes were defined, cyclic and non-cyclic, based on Spellmen et al. (1998) (see **Table S7**). For each SVM run, the full set of positive instances (cyclic expression) and negative instances (non-cyclic expression) was used to generate 100 balanced (i.e. 1-to-1 ratio of positive to negative) inputs to ensure that final evaluation is not biased by the fact that most of the genome it not cyclically expressed under any cell-cycle phase. Genes were only selected for the input of a SVM run if at least one TF-target or TF-TF interaction feature was present. In addition to the general cell-cycle model, an SVM model was established for predicting genes in each cell-cycle phase. Models were constructed as above expect that classes were defined as expression during a specific phase of the cell-cycle, again based on data from non-cyclic, based on Spellmen et al. (1998).

Each balanced input set was further divided for 10-fold cross validation with SVM implemented in Weka (Hall et al., 2009). Each run was optimized using a grid search of two parameters: (1) C: the minimum distance between the positive and negative groups, and (2) R: the ratio of negative to positive examples in the training set. The tested values of the two parameters were: C = (0.01, 0.1, 0.5, 1, 1.5, 2.0) and R = (0.25, 0.5, 1, 1.5, 2, 2.5, 3, 3.5, 4). For each pair of parameters, performance was measured using the prediction scores values averaged across the 100 balanced input sets and the reported AUC-ROC was calculated using this average score. For each choice of positive class and feature set, the pair of grid search parameters which maximized the average AUC-ROC was used to define the representative model for that predictor and calculate the reported AUC-ROC for that predictor.

### Evaluating the relationship between model performance, class and feature

The effect of the phase (general cell-cycle, G1, S, S/G2, G2/M or M/G1) of expression being predicted (class) and the data set (ChIP-Chip, Deletion, PWM, Expert PWM or PBM) from which TF-target interactions were derived (feature) on the performance of each SVM model was evaluated using analysis of variance (ANOVA). This was done using the “aov” function in the R statistical language using the following model:

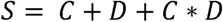

Where “S” is the representative AUC-ROC score of the SVM model, “C” is a categorical feature representing the positive-class set (cyclic expression or a specific phase of expression), and “D” is a categorical feature representing the data set of regulations used.

### Importance of features to predicting cell-cycle expression

The importance of a feature for each model was determined by rerunning each SVM model using the best pair of parameters with the options “-i -k” in order to generate an output files with class and features statistics. From the resulting output file, custom Python scripts were used to extract the weight value for each of the features used in the linear classifier. Features were then ordered by their weight to determine importance, such that the feature with the largest positive value (most strongly associated with the positive class) had the highest rank and the feature with the largest negative value (most strongly associated with the negative class) had the lowest rank. Because multiple features often had the same weight value, we defined cutoff scores for the 10th and 25th percentile conservatively, such that the cutoff for the Xth percentile of positive features was smallest weight above which includes X% or less of all features and the Xth percentile of negative features was the largest weight below which includes X% or less of all features. The effect of this is observed most prominently in the 25th percentile features sets as ties between feature weights were more common towards the middle of the weight distributions.

### GO Analysis

GO annotation for genes in *S. cerevisiae* were obtained from the Saccharomyces Genome Database (http://www.yeastgenome.org/download-data/curation). The significance of enrichment of a particular term in a set of important TF compared to the incidence of the GO annotation across the genome was determined using the Fisher’s Exact Test and adjusted for multiple-hypothesis testing using the Benjamini-Hochberg method (Benjamini and Hochberg, 1995).

## Acknowledgements

We would like to thank Zing Tsung-Yeh Tsai for his advice regarding approaches to modelling FFLs as well as Christina Azodi and Melissa Lehti-Shiu for reading this manuscript. This work was supported in part by grant from the National Science foundation (IOS-1546617, DEB- 1655386), and the DOE Great Lakes Bioenergy Research Center (DOE Office of Science BER DE-SC0018409) to S.-H.S.

## Supplemental Tables

**TableS1:** TF-target feature counts and target gene coverage by data set

**TableS2:** FFL feature counts and target gene coverage by data set

**Table S3:** AUR-ROC of ChIP TF-target interaction models on genes covered by ChIP FFLs

**Table S4:** Number of features in combined ChIP and Deletion models

**TableS5:** Importance of annotated cell-cycle regulations to prediction of different phases of expression in ChIP and Deletion data sets

**TableS6:** Enrichment of GO terms in TFs important to predicting phase of cell-cycle expression in ChIP

**Table S7:** Cell-cycle phase of S. cerevisiae genes

## References

Alon, Uri. An Introduction to Systems Biology. London: Chapman & Hall/CRC, 2007.

Alon U (2007) Network motifs: theory and experimental approaches. Nat Rev Genet 8: 450–461

Badis G, Berger MF, Philippakis AA, Talukder S, Gehrke AR, Jaeger SA, Chan ET, Metzler G, Vedenko A, Chen X, et al (2009) Diversity and complexity in DNA recognition by transcription factors. Science 324: 1720–1723

Bean JM, Siggia ED, Cross FR (2005) High functional overlap between MluI cell-cycle box binding factor and Swi4/6 cell-cycle box binding factor in the G1/S transcriptional program in Saccharomyces cerevisiae. Genetics 171: 49–61

Beer MA, Tavazoie S (2004) Predicting gene expression from sequence. Cell 117: 185–198

Benveniste D, Sonntag H-J, Sanguinetti G, Sproul D (2014) Transcription factor binding predicts histone modifications in human cell lines. Proc Natl Acad Sci U S A 111: 13367–13372

Berger MF, Bulyk ML (2009) Universal protein-binding microarrays for the comprehensive characterization of the DNA-binding specificities of transcription factors. Nat Protoc 4: 393–411

Bertoli C, Skotheim JM, de Bruin RAM (2013) Control of cell cycle transcription during G1 and S phases. Nat Rev Mol Cell Biol 14: 518–528

de Boer CG, Hughes TR (2012) YeTFaSCo: a database of evaluated yeast transcription factor sequence specificities. Nucleic Acids Res 40: D169–79

Buck MJ, Lieb JD (2004) ChIP-chip: considerations for the design, analysis, and application of genome-wide chromatin immunoprecipitation experiments. Genomics 83: 349–360

Bulyk ML (2007) Protein binding microarrays for the characterization of DNA-protein interactions. Adv Biochem Eng Biotechnol 104: 65–85

Cai L, Tu BP (2012) Driving the cell cycle through metabolism. Annu Rev Cell Dev Biol 28: 59–87

Chikina MD, Huttenhower C, Murphy CT, Troyanskaya OG (2009) Global prediction of tissue-specific gene expression and context-dependent gene networks in Caenorhabditis elegans. PLoS Comput Biol 5: e1000417

Csikász-Nagy A, Kapuy O, Tóth A, Pál C, Jensen LJ, Uhlmann F, Tyson JJ, Novák B (2009) Cell cycle regulation by feed-forward loops coupling transcription and phosphorylation. Mol Syst Biol 5: 236

van der Felden J, Weisser S, Brückner S, Lenz P, Mösch H-U (2014) The transcription factors Tec1 and Ste12 interact with coregulators Msa1 and Msa2 to activate adhesion and multicellular development. Mol Cell Biol 34: 2283–2293

Furey TS (2012) ChIP-seq and beyond: new and improved methodologies to detect and characterize protein-DNA interactions. Nat Rev Genet 13: 840–852

Garcia-Gimeno MA, Struhl K (2000) Aca1 and Aca2, ATF/CREB activators in Saccharomyces cerevisiae, are important for carbon source utilization but not the response to stress. Mol Cell Biol 20: 4340–4349

Hall M, Frank E, Holmes G, Pfahringer B, Reutemann P, Witten IH (2009). The WEKA Data Mining Software: An Update. SIGKDD Explorations, 11

Hanlon SE, Rizzo JM, Tatomer DC, Lieb JD, Buck MJ (2011) The stress response factors Yap6, Cin5, Phd1, and Skn7 direct targeting of the conserved co-repressor Tup1-Ssn6 in S. cerevisiae. PLoS One 6:e19060

Harbison CT, Gordon DB, Lee TI, Rinaldi NJ, Macisaac KD, Danford TW, Hannett NM, Tagne J-B, Reynolds DB, Yoo J, et al (2004) Transcriptional regulatory code of a eukaryotic genome. Nature 431: 99–104

Hauser M, Narita V, Donhardt AM, Naider F, Becker JM (2001) Multiplicity and regulation of genes encoding peptide transporters in Saccharomyces cerevisiae. Mol Membr Biol 18:105–12

Iyer VR, Horak CE, Scafe CS, Botstein D, Snyder M, Brown PO (2001) Genomic binding sites of the yeast cell-cycle transcription factors SBF and MBF. Nature 409: 533–538

Jarolim S, Ayer A, Pillay B, Gee AC, Phrakaysone A, Perrone GG, Breitenbach M, Dawes IW (2013) Saccharomyces cerevisiae genes involved in survival of heat shock. G3 3: 2321–2333

Jolma A, Yin Y, Nitta KR, Dave K, Popov A, Taipale M, Enge M, Kivioja T, Morgunova E, Taipale J (2015) DNA-dependent formation of transcription factor pairs alters their binding specificity. Nature 527: 384–388

Jorgensen P, Nishikawa JL, Breitkreutz B-J, Tyers M (2002) Systematic identification of pathways that couple cell growth and division in yeast. Science 297: 395–400

Juven-Gershon T, Hsu J-Y, Theisen JW, Kadonaga JT (2008) The RNA polymerase II core promoter - the gateway to transcription. Curr Opin Cell Biol 20: 253–259

Kazemian M, Pham H, Wolfe SA, Brodsky MH, Sinha S (2013) Widespread evidence of cooperative DNA binding by transcription factors in Drosophila development. Nucleic Acids Res 41: 8237–8252

Koranda M, Schleiffer A, Endler L, Ammerer G (2000) Forkhead-like transcription factors recruit Ndd1 to the chromatin of G2/M-specific promoters. Nature 406: 94–98

Lambert JR, Bilanchone VW, Cumsky MG (1994) The ORD1 gene encodes a transcription factor involved in oxygen regulation and is identical to IXR1, a gene that confers cisplatin sensitivity to Saccharomyces cerevisiae. Proc Natl Acad Sci U S A 91:7345–9

Lelli KM, Slattery M, Mann RS (2012) Disentangling the many layers of eukaryotic transcriptional regulation. Annu Rev Genet 46: 43–68

Li M, Hada A, Sen P, Olufemi L, Hall MA, Smith BY, Forth S, McKnight JN, Patel A, Bowman GD, et al (2015a) Dynamic regulation of transcription factors by nucleosome remodeling. Elife. doi: 10.7554/eLife.06249

Li Y, Chen C-Y, Kaye AM, Wasserman WW (2015b) The identification of cis-regulatory elements: A review from a machine learning perspective. Biosystems 138: 6–17

Macneil LT, Walhout AJM (2011) Gene regulatory networks and the role of robustness and stochasticity in the control of gene expression. Genome Res 21: 645–657

Mannhaupt G, Schnall R, Karpov V, Vetter I, Feldmann H (1999) Rpn4p acts as a transcription factor by binding to PACE, a nonamer box found upstream of 26S proteasomal and other genes in yeast. FEBS Lett 450: 27–34

Marion RM, Regev A, Segal E, Barash Y, Koller D, Friedman N, O’Shea EK (2004) Sfp1 is a stress- and nutrient-sensitive regulator of ribosomal protein gene expression. Proc Natl Acad Sci U S A 101: 14315–14322

Martinez-Pastor MT, Marchler G, Schuller C, Marchler-Bauer A, Ruis H, Estruch F (1996) The Saccharomyces cerevisiae zinc finger proteins Msn2p and Msn4p are required for transcriptional induction through the stress response element (STRE). EMBO J 15: 2227–35.

McInerny CJ, Partridge JF, Mikesell GE, Creemer DP, Breeden LL (1997) A novel Mcm1-dependent element in the SWI4, CLN3, CDC6, and CDC47 promoters activates M/G1-specific transcription. Genes Dev 11: 1277–1288

Miller JA, Widom J (2003) Collaborative competition mechanism for gene activation in vivo. Mol Cell Biol 23: 1623–1632

Ng DT, Spear ED, Walter P (2000) The unfolded protein response regulates multiple aspects of secretory and membrane protein biogenesis and endoplasmic reticulum quality control. J Cell Biol 150: 77–88

O’Malley RC, Huang S-SC, Song L, Lewsey MG, Bartlett A, Nery JR, Galli M, Gallavotti A, Ecker JR (2016) Cistrome and Epicistrome Features Shape the Regulatory DNA Landscape. Cell 165: 1280–1292

Panchy N, Wu G, Newton L, Tsai C-H, Chen J, Benning C, Farré EM, Shiu S-H (2014) Prevalence, evolution, and cis-regulation of diel transcription in Chlamydomonas reinhardtii. G3 4: 2461–2471

Pett JP, Korenčič A, Wesener F, Kramer A, Herzel H (2016) Feedback Loops of the Mammalian Circadian Clock Constitute Repressilator. PLoS Comput Biol 12: e1005266

Reimand J, Vaquerizas JM, Todd AE, Vilo J, Luscombe NM (2010) Comprehensive reanalysis of transcription factor knockout expression data in Saccharomyces cerevisiae reveals many new targets. Nucleic Acids Res 38: 4768–4777

Schmitt AP, McEntee K (1996) Msn2p, a zinc finger DNA-binding protein, is the transcriptional activator of the multistress response in Saccharomyces cerevisiae. Proc Natl Acad Sci U S A 93: 5777–82

Segal E, Raveh-Sadka T, Schroeder M, Unnerstall U, Gaul U (2008) Predicting expression patterns from regulatory sequence in Drosophila segmentation. Nature 451: 535–540

Spellman PT, Sherlock G, Zhang MQ, Iyer VR, Anders K, Eisen MB, Brown PO, Botstein D, Futcher B (1998) Comprehensive identification of cell cycle-regulated genes of the yeast Saccharomyces cerevisiae by microarray hybridization. Mol Biol Cell 9: 3273–3297

Spitz F, Furlong EEM (2012) Transcription factors: from enhancer binding to developmental control. Nat Rev Genet 13: 613–626

Spoko R, Huang D, Preston N, Chua G, Papp B, Kafadar K, Synder M, Oliver SG, Cyret M, Hughes TR, Boone C, Andrews B (2006) Mapping pathways and phenotypes by systematic gene overexpression. Mol Cell 21:319–30.

Stormo GD, Schneider TD, Gold L, Ehrenfeucht A (1982) Use of the “Perceptron” algorithm to distinguish translational initiation sites in E. coli. Nucleic Acids Res 10: 2997–3011

Tomancak P, Beaton A, Weiszmann R, Kwan E, Shu S, Lewis SE, Richards S, Ashburner M, Hartenstein V, Celniker SE, et al (2002) Systematic determination of patterns of gene expression during Drosophila embryogenesis. Genome Biol 3: RESEARCH0088

Uygun S, Seddon AE, Azodi CB, Shiu S-H (2017) Predictive Models of Spatial Transcriptional Response to High Salinity. Plant Physiol 174: 450–464

Vizoso-Vazquez A, Lamas-Maceiras M, Becerra M, Gonzalez-Siso MI, Rodriquez-Belmonte E, Cerdan ME (2012) Ixr1p and the control of the Saccharomyces cerevisiae hypoxic response. Appl Microbiol Biotechnol 94:173–84

Wasserman WW, Sandelin A (2004) Applied bioinformatics for the identification of regulatory elements. Nat Rev Genet 5: 276–287

White SE, Riles L, Cohen BA (2009) A systematic screen for transcriptional regulators of the yeast cell cycle. Genetics 181:435–46

Wittenberg C, Reed SI (2005) Cell cycle-dependent transcription in yeast: promoters, transcription factors, and transcriptomes. Oncogene 24: 2746–2755

Xie Y, Varshavsky A (2001) RPN4 is a ligand, substrate, and transcriptional regulator of the 26S proteasome: a negative feedback circuit. Proc Natl Acad Sci U S A 98: 3056–3061

Xu Z, Norris D (1998) The SFP1 gene product of Saccharomyces cerevisiae regulates G2/M transitions during the mitotic cell cycle and DNA-damage response. Genetics 150: 1419–1428

Zhu C, Byers KJRP, McCord RP, Shi Z, Berger MF, Newburger DE, Saulrieta K, Smith Z, Shah MV, Radhakrishnan M, et al (2009) High-resolution DNA-binding specificity analysis of yeast transcription factors. Genome Res 19: 556–566

Zhu G, Spellman PT, Volpe T, Brown PO, Botstein D, Davis TN, Futcher B (2000) Two yeast forkhead genes regulate the cell cycle and pseudohyphal growth. Nature 406: 90–94

Zou C, Sun K, Mackaluso JD, Seddon AE, Jin R, Thomashow MF, Shiu S-H (2011) Cis-regulatory code of stress-responsive transcription in Arabidopsis thaliana. Proc Natl Acad Sci U S A 108: 14992–14997

